# A Postdoctoral Training Program in Bioimage Analysis

**DOI:** 10.1101/2024.05.13.593910

**Authors:** Beth A Cimini, Callum Tromans-Coia, David Stirling, Suganya Sivagurunathan, Rebecca Senft, Pearl Ryder, Esteban Miglietta, Paula Llanos, Nasim Jamali, Barbara Diaz-Rohrer, Shatavisha Dasgupta, Mario Cruz, Erin Weisbart, Anne E Carpenter

## Abstract

We herein describe a postdoctoral training program designed to train biologists with microscopy experience in bioimage analysis. We detail the rationale behind the program, the various components of the training program, and outcomes in terms of works produced and the career effects on past participants. We analyze the results of an anonymous survey distributed to past and present participants, indicating overall high value of all 12 rated aspects of the program, but significant heterogeneity in which aspects were most important to each participant. Finally, we propose this model as a template for other programs which may want to train experts in professional skill sets, and discuss the important considerations when running such a program. We believe that such programs can have extremely positive impact for both the trainees themselves and the broader scientific community.

## Introduction

The past 10 years have seen a veritable explosion of interest in the quantitative analysis of microscopy images and the development of software for this purpose. The pioneering image analysis programs in this field, such as ImageJ^1^/Fiji^2^, CellProfiler^3^, and ilastik^4^, have been joined by an ever-expanding array of helpful tools, libraries, and plugins. As of this writing, there are 63 participating software packages and 6 community organizations (such as societies) active on the Scientific Community Image Forum (forum.image.sc)^5^. More than ∼80 image analysis software packages or languages were listed in a recent review^6^.

Unfortunately, this upward trend in the development of new image analysis software has not meant universal adoption and utilization. Despite the efforts of groups such as the Network of European Union Biomage Analysts (NEUBIAS^7^), the Global Bioimage Analysts’ Society (GloBIAS), AI4Life, and the Royal Microscopical Society (RMS), progress in the community’s ability to effectively use these tools has been uneven. Findings from a recent survey indicate that while AI and machine learning (ML) approaches to image analysis are viewed as potentially transformative, access to analysis can be a major barrier^8^. Global surveys^9,10^ show that computational comfort is not always high among the users of bioimage analysis software, and tends to be lower in users who spend more of their time acquiring rather than analyzing the images.

The last several years have generally seen an increase in what Global Bioimaging refers to as imaging scientists, professional scientists typically found in core facilities who can guide colleagues through creating successful microscopy experiments^11^ . In parallel, the increasing need for computational image analysis expertise has created a relatively new career path - that of the bioimage analyst^12–14^. Bioimage analysts can serve as a bridge between disciplines including biology, computer science, bioinformatics, and physics that may have common techniques and/or goals but different languages and emphases. Bioimage analysts interact with biologists who have images to analyze but possess limited computational understanding, as well as with computer scientists with advanced image processing skills but no biological domain knowledge. There are successful bioimage analysts from many backgrounds, including but not limited to biology, medicine, physics, and computer science.

Having observed Jennifer Waters’ successful training program for microscopy core facility managers (started in 2013 at Harvard Medical School^15^), we (BAC/AEC) realized how valuable a parallel image analysis focused program would be to meet the growing needs for bioimage analysts in academia and industry. In 2019, we launched what we believe was the first-ever Bioimage Analysis Postdoctoral Training Program with two openings at the Broad Institute.

Since then, seven scientists have completed the program, with four more in progress. We herein describe the components of the program, attempt to quantify the most important aspects, and provide suggestions for others wishing to replicate this model.

## Results

### Applicant pool

The goal of our program is to train PhD-level researchers who have deep experience in biological experimentation, including microscopy, and are motivated to develop image analysis expertise over the course of 2-3 years. The training largely follows an apprenticeship model; solving various image analysis tasks forms the core of the training experience. In six calls as of this writing, >200 total candidates applied to the program. The past and current program members (n=11) have included 7 women and 4 men from 8 countries of origin on 4 continents; all had obtained a PhD, with two also having earned MDs. This broad and deep applicant pool, even during both a pandemic and a so-called “postdoc crisis”^16,17^, lends support to anecdotal reports of increased interest in “alternate” (non-tenure-track-faculty) careers.^18,19^

Successful applicants to the program were all skilled in creating bioimages and expressed enjoyment in collaborating with colleagues from diverse disciplinary backgrounds, but possessed varying levels of previous computational experience at the time of their application. In general, accepted program members fell into two broad computational experience groups: (i) those with little to no computational experience but with a passion to learn and broadly improve these skills, and (ii) those with moderate (but typically self-taught) computational experience on a limited range of biological images who sought to work on more diverse data and to professionalize their skills by working alongside formally-trained computer scientists.

### Program structure

Scientists entering the program customize a template curriculum (Supplementary Data 1) which supports development in a variety of areas, including computational skills, collaboration, project management, and outreach. In the early months of the program, all program members focus on specific tasks such as reviewing existing tutorial materials, becoming familiar with version control systems and with cloud computing, and observing and conducting “office hour” sessions with other researchers seeking bioimage analysis assistance; these tasks ensure they attain the broad training base needed for successful completion of the program. Because backgrounds and eventual career goals vary widely, after developing a well-rounded initial knowledge base, the members are encouraged to specialize and the recommendations are tailored accordingly. Quarterly career chats, including an annual performance review process, ensure periodic evaluation of big-picture development and progression of skills towards eventual goals.

In addition to emphasizing peer collaboration and joint problem solving, interaction with longer-term members of the lab is a key aspect of the training program. In addition to direct training from the lab Principal Investigator, the program is embedded in a lab with several software engineers focused on developing bioimage analysis tools and an experienced image analyst. In addition to the typical knowledge transfer that occurs in a scientific laboratory through both informal interactions encouraged by proximity and formal lab meeting presentations, postdocs, staff, and PI all participate in a 2-hour weekly group check-in meeting wherein all attendees share in a casual manner challenges, successes, or learning opportunities from the previous week. Trainees not only receive direct feedback during these check-ins but also witness staff develop and troubleshoot advanced methods and workflows. The staff members also make the whole program more sustainable over time by serving as a repository of knowledge and experience.

Having many robust sources of institutional knowledge is critical in such a training program, which in our experience is supported by text-based troubleshooting channels (such as Slack and a private StackOverflow instance) which are private to the outside world (removing some anxiety around asking even “basic” questions) but public to the group, meaning questions and answers are searchable by current and future members since similar problems often recur. Lab employees of all seniority levels are encouraged to answer but also to ask questions in these channels, further emphasizing to training program members that in a multidisciplinary environment, no single person knows all answers and learning and development are critical at all career stages.

While the emphasis on particular development areas varies by program member, all members in general work on a variety of collaborative image analysis projects, which span from a few hours in a single day to hundreds of hours over months or years. Each member works on approximately 10 projects per year. Collaborations are drawn from both academia and industry, and may be funded by the collaborating lab or by the program’s own grants. Where externally funded, program members are typically required to track the number of hours spent on the project, as collaborators are billed based on hours spent. This process allows members to hone project- and time-management skills which are rarely taught in academic settings but highly sought by employers. Additional exposure to diverse image analysis tasks and collaborator communication comes from conducting open “office hours” with scientists from across the globe, as well as by answering questions on the online Scientific Community Image Forum (forum.image.sc). Presentation and educational skills are practiced by conducting live workshops (in-person or online), writing blog posts, and/or recording video tutorials. Most program members write at least one co-first or first-author paper while in the program, including surveys of existing tools or open needs, papers detailing protocols developed during their tenure, or reports on new or improved software created.

### Community impact

Beyond training individual scientists, the program benefits the broader bioimaging community. In less than five years of operation, scientists in the training program have collaborated with dozens of labs, often training many collaborators within them, have first- or co-first authored nine publications (with a collective 871 citations) and earned middle authorship on 9 more publications (with collectively 185 citations). The companion tutorial website to one of these first author publications has received >4,000 unique visitors (with >50 unique visitors in 15 countries) and has been translated into three additional languages, with six more in progress.

Program members led or assisted in 29 workshops, reaching >1200 participants either in person or on Zoom (on at least four continents), and were responsible for writing and/or translating eight CellProfiler tutorial exercises. They created 17 tutorial videos with >38,000 collective views on YouTube, and wrote 17 blog posts which received >16,000 page views. They conducted 118 office hours and posted collectively >2,100 times on the Scientific Community Image Forum (forum.image.sc). These metrics highlight the impact of the program on the wider bioimaging community beyond the individual participants and specific collaborating labs.

### Program participant survey

To assess the effects of our past efforts and better target future efforts, all 11 program participants were anonymously surveyed in the fall of 2023 (see also Materials and Methods); 9 participants (5/7 program alumni, 4/4 current members) responded. Respondents were asked to score (on a scale of 1-7) and rank (1st to 12th) the importance of 12 aspects of the program (Figure 1). In general, most activities were highly scored, with all activities in all subgroups receiving a mean >4 (on a scale of 1 to 7). Working on individual image analysis projects and gaining experience in coding received unanimous 7/7 scores from all participants, with answering forum questions, gaining data science experience, and communication skills all additionally receiving a mean score >6 among both current and past participants.

**Figure 1:**
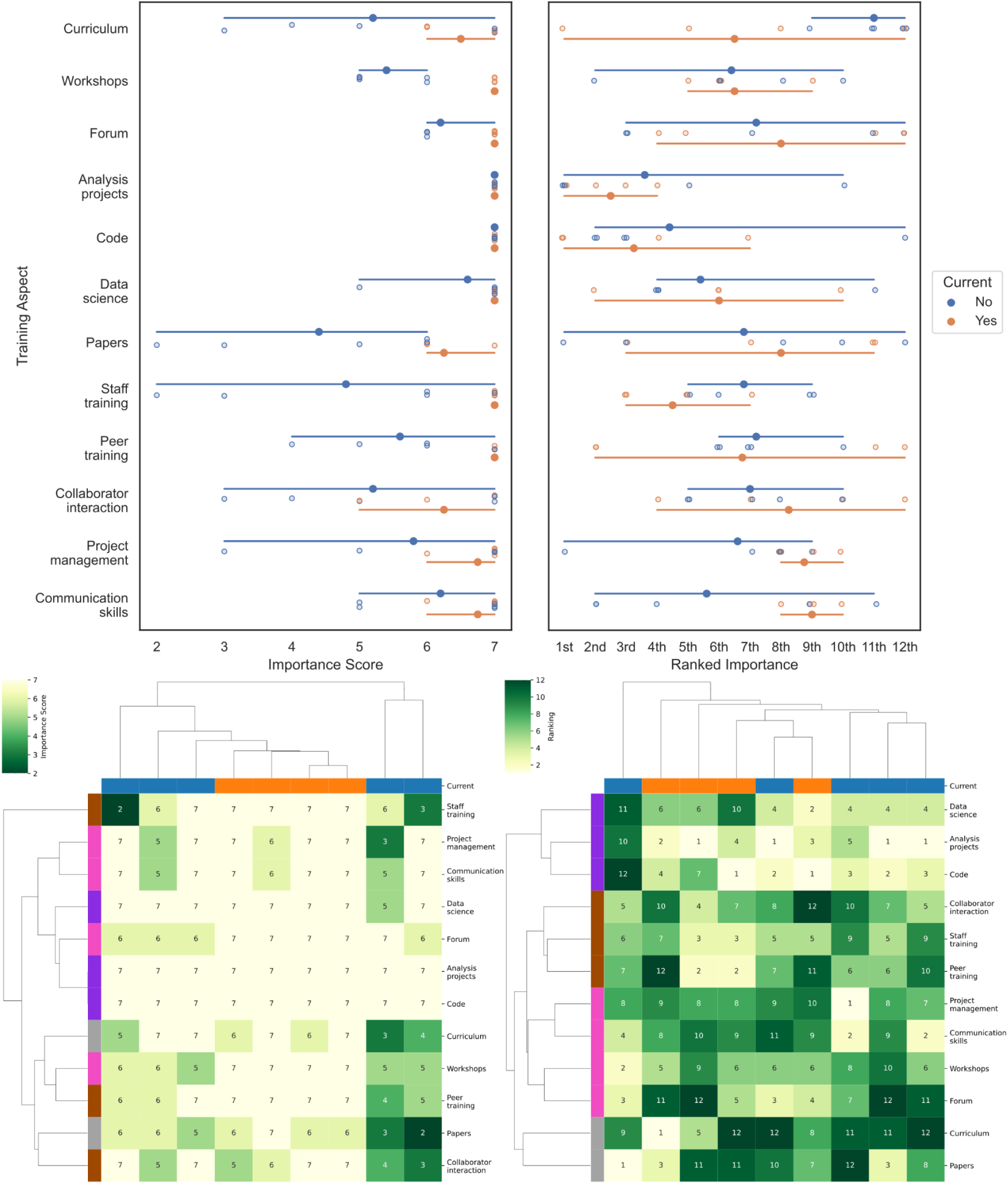
Relative scores (left) and rankings (right) of the value of 12 aspects of the training program to the participant’s training experience. Aspects include: having an official training curriculum, running workshops, working with the image.sc forum, working on individual analysis projects, writing code, working on data-science-heavy-projects, writing papers, training with/from permanent staff members (image analyst/software engineers), training with/from other postdoc program members, interaction with collaborators, explicit and/or implicit training on project management skills, and explicit and/or implicit training on communication skills. Scores were on a scale of 1-7; rankings were performed 1st-12th with no possibility of ties. Top: Score results are shown broken down by past vs current members. Bottom: Results are shown hierarchically clustered. Columns are colored by whether the answers come from a current member; rows are colored by thematic axes derived from the clustering of the rankings.

Analysis of the rankings revealed more diversity among the respondents: with the exception of “training with staff”, all 12 program aspects had at least one vote for being in the top three most important aspects and at least one vote in the bottom three least important aspects (Figure 1). Among the largest disparities in rankings between current and past members was the higher importance placed on the curriculum by current members and higher importance placed on communication skills and project management skills by past members. These results highlight the importance of consideration of a growth-supporting training experience when designing such a program; creating structure and clear expectations is important to engender trust and safety in initial stages but flexibility and widely transferable job skills become increasingly more valued in later stages. Clustering analysis shows broadly similar response patterns across the majority of participants (Figure 1). To look for themes across the 12 aspects, we further examined the row clustering of the aspect rankings (which were far more varied than the scores), four approximate “thematic axes” are created by clustering, which could be loosely termed “hard skills” (analysis projects, code, and data science), “co-training” (interaction with peers, staff, and collaborators) “outreach and soft skills” (workshops, forum participation, communication, and project management), and “other” (the formal curriculum and writing papers).

We also asked program members about their preferred program length (Figure 2); the modal answer was 18-24 months. The alumni rated the program length to be “about right”, both in terms of their training experience, as well as the impact of the program on their personal lives (e.g. salary, location requirement). Asked if joining a training program was the right choice for them, of the five past members who completed the survey, four answered “Yes” (or a variant thereon) and one answered “Maybe”. Both current and past members were also asked to reflect on their career goals (Figure 2).

**Figure 2:**
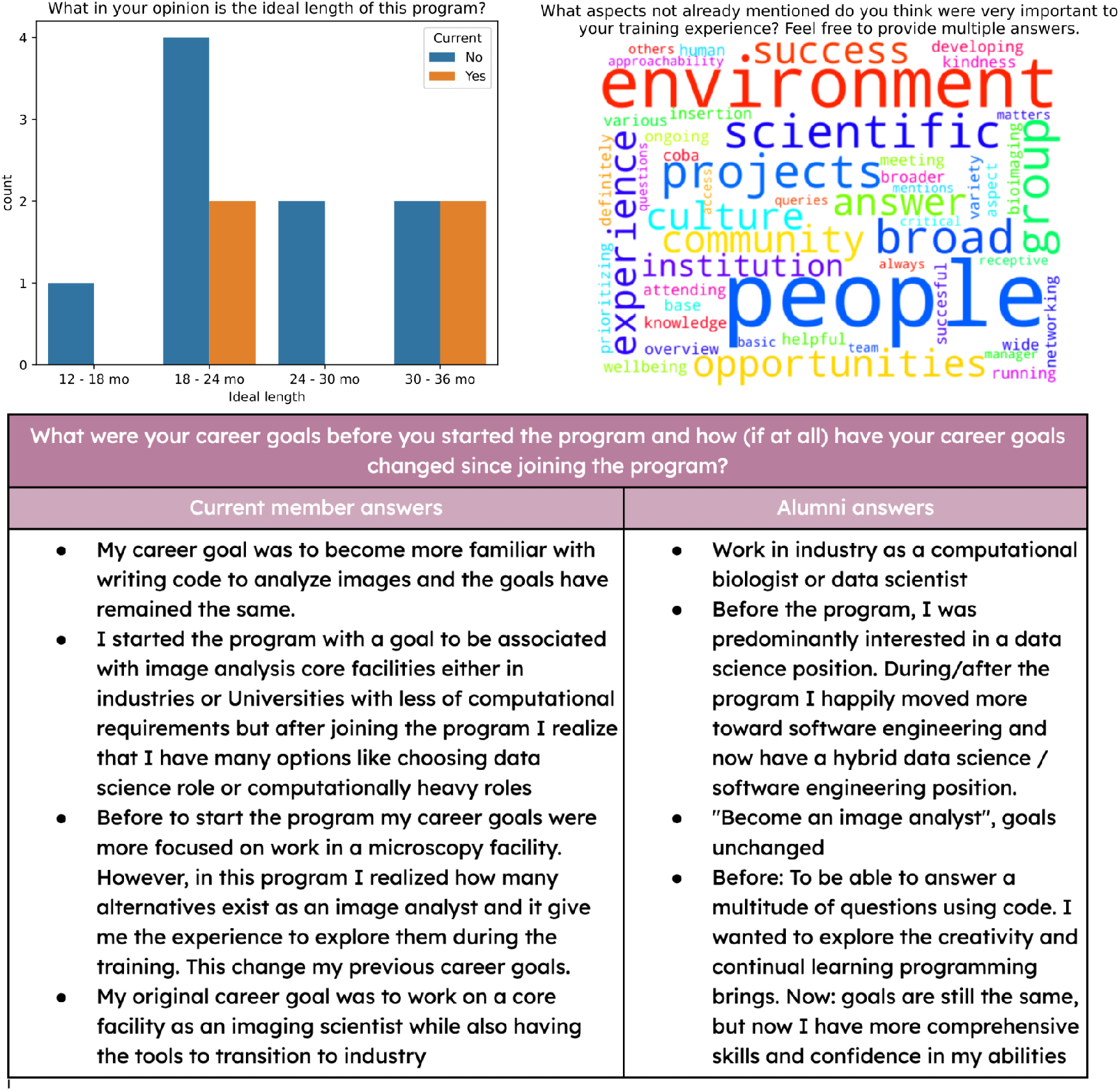
Selected responses to other survey questions. Top left: respondent feelings on the ideal program length (multiple answers could be chosen by each respondent). Top right: wordcloud of other factors. Bottom: all responses to the question: “What were your career goals before you started the program and how (if at all) did they change during or since the program?”

Responses to the question “What aspects not already mentioned do you think were very important to your training experience?” highlighted a largely-intangible aspect to be important for success: a welcoming and supportive environment for learning and a culture that emphasizes growth and development. Responses to “What aspects do you think were missing from your training experience?” tended to be for minor specific improvements (see Supplementary Data 2); common themes included 1) a desire in some participants for more interaction with image analysts outside of their own lab and 2) an interest in continued participation in biological projects after the completion of the initial image analysis, either through performing additional data analysis and/or hearing from the collaborators how the results supported or refuted their research hypothesis. Responses to a final free-form question (see Supplementary Data 2) highlighted the importance of the program in spreading knowledge of open-source solutions to the broader scientific community and the importance of practical experience in acquiring advanced image analysis skills.

### Trainee impact

While trainees learn a number of skills while in the program (including software engineering skills, experience with large scale workflows, version control systems, and project management), the most critical outcome of any training program is the ultimate impact on trainee careers. Through a diversity of projects and collaborators, our program helps trainees realize the breadth of careers available in and beyond bioimage analysis. The program also has implemented lab events such as an annual ‘career day’ where lab members dedicate time to work on enhancing their CVs and to get feedback from others in the group, including lab principal investigators. Practice talks for interviews with feedback from lab members are also encouraged, another practice that creates a culture benefitting trainees. As the number of program participants grows over time, another emergent effect of the program is creating a network of past postdocs who have pursued various career paths and can advise current participants on selecting and pursuing next steps. In practice, this has ranged from career discussions to interview prep and advice on negotiating an offer. Careers of program participants include scientist and data scientist positions at startups, biotechs, and pharmaceutical companies of varying sizes; advanced postdoctoral positions, software engineer and analyst positions, and founding roles in university image analysis core groups. The diversity of these positions indicates the broad usefulness of skills gained in the course of the program.

The support of the program for pursuing different careers fosters an open culture where participants can easily explore their options and ultimately realize their desired career path and gain the skills and experience to successfully pursue it.

## Discussion

We find an apprenticeship-style postdoctoral training program in bioimage analysis to be successful, benefitting both the participants and the broader community. Participants in the program gained valuable experience and skills for future careers, and collectively helped thousands of other researchers through the software and educational materials they created. While our program focuses on training scientists with a previous background in bioimaging to acquire additional image analysis skills, other programs, such as at the Center for Imaging Technology and Education at Harvard Medical School and the Advanced Imaging Center at HHMI-Janelia, focus on bioimaging more broadly. One could envision similar programs being created for other collaborative biotechnical disciplines, each with its own specialization in terms of which skills are already possessed by entering trainees and which are developed during the program.

Despite the catalytic impact, running an apprenticeship-style program is not without its challenges. In a typical core-facility(-like) environment where projects are funded by charging collaborators for increments of time, there can be conflicts in the interests of the trainee and the facility. Trainees benefit from significant amounts of unstructured, unbilled learning time during problem solving, and from departing the program as soon as “sufficient skills’’ are acquired. By contrast, the facility’s financial health and scientific sustainability is threatened if insufficient hours are billed and program members depart before contributing to teaching newer members. These tensions also exist for conventional staff roles in facilities (especially computational facilities)^13,20^, but will of course be higher the more “unbillable” time a given trainee is allotted. These tensions could be reduced by creating funding opportunities for training programs that cover fractions of trainee and/or mentor salaries, which would reduce the need to fully recover salary costs. Funders of such programs could also require explicit training standards that would ensure that a promised “training role” is not simply a standard staff role with lower salary. While we are not aware of any such opportunities currently, we hope that the success of programs like ours may inspire funders to create such calls or to otherwise experiment with creative solutions that simultaneously train much-needed technical experts while providing employment for promising scientific trainees. Our program has only been financially feasible due to grants from the National Institutes of Health supporting the CellProfiler and ImageJ software projects, which included funding to support outreach and training as well as some collaborations; expansion of these kinds of grants would be catalytic in strengthening the workforce in valuable computational areas.

In our experience, the aspect most crucial for the success of this program has been the selection of participants with strong communication and organizational skills, who are willing to persist with enthusiasm even while encountering the frustrations always present when mastering difficult new techniques. Equally important, we believe, has been our intentional cultivation of a supportive environment that promotes collaboration and encourages the attitude of “I don’t know, but I’m interested to find out”. Program members report comfort asking questions and advocating for their personal and professional needs without the fear of judgment, and participate in an annual lab retrospective where they can anonymously suggest changes to policies and procedures. Creation of protected time periods for reading and learning and conscious focus on acquiring skills and tracking progress towards stated goals (such as during periodic “career chats”) are also critical for creating a safe environment for training.

As one program member nicely summarized: “I think the most challenging aspect of this program is balancing providing enough framework and structure to help individuals find their career path without making a curriculum that is more rigid than useful. I think the program benefits enormously when the leadership, alumni, and senior postdocs help to provide a framework for incoming postdocs to understand what skills they should develop to fit the career paths that interest them.” Balancing the various factors required to create a fiscally sustainable but successful program is an undeniably difficult task, but the impact on individual scientists and the broader community they go on to serve, is enormous.

## Materials and Methods

An anonymous Google Form was distributed by email to all current and former members of the program, with responses received from 5/7 alumni and 4/4 current members. Answers to questions in the first section (relative importance of various activities) were not randomized in order to be able to detect patterns of similar responses. Answers to questions in the second section (program length and career goals) were subgrouped by alumni status but otherwise randomized, while free-text answers in the third section were fully randomized. A copy of the survey is available as Supplementary Data 3 and the answers to all questions but the final question (for feedback the user did not wish to be included in the paper) is available as Supplementary Data 2.

Graphs were run in Python 3.10 using the Jupyter^21^, pandas^22^, seaborn^23^, and wordcloud^24^ packages; source data and code are available at https://github.com/ciminilab/2024_PostdocProgram.

## Acknowledgements

We gratefully acknowledge the contributions of the members of the Carpenter lab (2019-2021) and Carpenter-Singh and Cimini labs (2021-present) in creating a welcoming and dynamic scientific training environment in which to host this program. Lab members are also thanked for their inputs on this manuscript. Members of the training program are listed in reverse-alphabetical authorship order, with no other ranking implied.

