## Supplementary Data 1 for "A Postdoctoral Training Program in Bioimage Analysis"

### Welcome

First of all, welcome to the Cimini lab Image Analysis team, and to the whole Imaging Platform! We're really excited to have you here, and you've been selected because we strongly believe that you have what it takes to succeed.

This is probably your first "dry" lab experience - it takes some getting used to! Your time will be mostly unstructured, as external events like incubation times and experimental plans don't give you a natural idea of when to do what. So, you will need to work out what works best for you to stay motivated on projects, decide when to switch priorities/tasks, and so on. Don't hesitate to ask the group for tips, as we tend to be fans of various project management/to-do tools. On the up side, you will find your life is less dominated by cells that won't cooperate, crummy antibodies, timepoints that interrupt your personal life, etc.

You will be exposed to such a breadth of projects here! Many of them are translational: many PIs have MD PhDs and see patients while also overseeing research. Our collaborators tend to be just amazing scientists and it's a delight to have our efforts going towards accelerating their work. If there are particular types of projects you're interested in, speak up- we can make sure you get first chance at those when they come in, or can think if there's a collaboration that makes sense to start or approach.

### Goals

The goal of this training program is to grow your skill set so that you can find gainful and fulfilling employment working to analyze biological images (or something in a related field). By the time you leave the lab, you should have all the skills needed for success:

- Broad expertise in image analysis software and a variety of biological problems
- Ability to dive in and enthusiastically explore new technologies, as this field constantly changes
- Strong organization and time management skills- working independently to manage your own time, efficiently juggling multiple tasks, setting and managing expectations, EXCELLENT documentation skills, and predicting, sticking to, and tracking a per-hour budget per project
- Strong communication skills, including one-on-one communication with collaborators, the ability to convey computational concepts to large groups of biologists (through talks or workshops), and the ability to communicate with software developers to convey bugs or feature requests

### Your long-term career ideas

- <what options are you open to at this time?>

### Roadmap

With that in mind, here is a roadmap to what you can expect in your time in the lab- while everyone's journey will be slightly different, this is how we envision your progression through the program, and how we'll make sure you are developing the skills we think you need to succeed!

- Months 0-3
  - Orientation to the basics with our administrator (email/calendar accounts, where to find information, etc)
  - Orientation to project management: project tracking, data locations, etc
    - Onboarding sections of these two documents
  - Orientation to CellProfiler- complete tutorials, workshop material, work on forum questions, etc
    - Please update materials as you go to make them even easier for beginners!
    - Suggested order - Translocation, Advanced Segmentation, 3D, Pixel Classification, QC
  - Work on initially assigned image analysis projects (start from week 2-3)
  - Track what you're doing on the Image Analysis Trello board
  - Start thinking about training area(s) of specialization. Previous areas have included (yours can be one of these or something totally different!):
    - Neuroscience projects
    - Time-lapse analysis projects
    - Imaging flow cytometry projects
    - Tissue projects
    - CellProfiler software engineering
    - Deep learning
    - Projects with major data-science components (Cell Painting/profiling)
    - Training and outreach
  - Attend at least one CellProfiler workshop to observe teaching styles and strategies
  - Read the Image.sc forum at least 1 hr/week (feel free to answer questions where you can)
  - Attend and observe office hour sessions
  - Please read the COBA Summary Document

    - If you don't already have tools you like for to-do list tracking, whiteboard, etc, check out the most recent project organization tool list
- Months 3-6
    - Become independent on individual image analysis projects, one at a time
        - Note: “independent” does not mean you never ask questions! No matter how skilled, we always ask each other for input on projects.
    - Explore other open source software packages- FIJI, Icy, ilastik, KNIME, Orbit, etc
    - Learn how to use cloud computing resources (AWS) responsibly
    - Outline discrete goals for area(s) of specialization
        - Asking Beth to be assigned to a specific type of project
        - Taking a class/attending seminars
        - Choosing a particular lab chore that aligns with a goal (ie database management or bash scripting)
        - Identifying alternate mentors inside or outside the group to help achieve goals
    - Assist at least one CellProfiler workshop
    - Answer questions on the Image.sc forum at least 1 hr/week
    - Co-lead office hour sessions
    - Begin using our time tracking software
- Months 6-12
    - Become independent on balancing multiple simultaneous image analysis projects
    - Continue exploring other open source software packages
    - Identify and attend a conference in the field
    - Write a blog post (see past ones as examples)
    - Set and work towards anniversary goal(s) for area(s) of specialization - check in with Beth (and any alternate mentors) monthly
    - Present at least one CellProfiler workshop to practice education and public speaking
    - Answer questions on the Image.sc forum at least 1 hr/week
    - Independently lead office hour sessions
- Months 12-24
    - Continue to work independently on multiple image analysis projects
    - Set and work towards year 2 goals for area(s) of specialization - check in with Beth (and any alternate mentors) monthly
        - Consider if you want an additional field for year 2!
    - Review job postings in the field every month to see what is out there, what skills they are looking for, and what skills you may want to think about developing further
        - If you must be very picky due to geography or other considerations, let us know if you see a position opening that is worth leaving the program early for
    - Present at a conference in the field
- Months 24+

- Continue to work independently on multiple image analysis projects
- Set and work towards any final goals for area(s) of specialization - check in with Beth (and any alternate mentors) monthly
- Identify future opportunities outside the lab and ask Anne and Beth for contacts that would facilitate your career path
