## Supplementary Data 3 for "A Postdoctoral Training Program in Bioimage Analysis"

### Postdoctoral Training Program Feedback form

This form has 3 sections, but should only take a few minutes depending on how much written open-ended feedback you want to provide (I am grateful for any and all but you're all busy people with busy lives!). Only the ~20 multiple choice questions are required, all open-ended questions are optional.

- First section on the importance of various aspects of the training
- Second section on program length and career goals - questions will be slightly different for past and present postdocs
- Third section for open ended responses about anything else

Responses will not go directly to me but will instead go to Erin for randomization of sections 2 and 3 before I see them, so while I hope you always feel comfortable giving me critical/negative feedback directly, in case you want to do so fully anonymously, that will be possible (assuming you don't out yourself with your name or a project only you worked on, but that will only out you for that one question). If you want to give me private feedback directly (my eyes only), you can do it here.

Not shared

* Indicates required question

Are you still in the program? *

- ○ Yes
- ○ No

How important do you think having an official training curriculum was/is? *

| | 1 | 2 | 3 | 4 | 5 | 6 | 7 | |
|---|---|---|---|---|---|---|---|---|
| Not at all important | ○ | ○ | ○ | ○ | ○ | ○ | ○ | Extremely important |

How valuable is/was running workshops to your training experience? *

| | 1 | 2 | 3 | 4 | 5 | 6 | 7 | |
|---|---|---|---|---|---|---|---|---|
| Not at all valuable | ○ | ○ | ○ | ○ | ○ | ○ | ○ | Extremely valuable |

How valuable is/was working with the image.sc forum to your training experience? *

| | 1 | 2 | 3 | 4 | 5 | 6 | 7 | |
|---|---|---|---|---|---|---|---|---|
| Not at all valuable | ○ | ○ | ○ | ○ | ○ | ○ | ○ | Extremely valuable |

How valuable is/was working on individual analysis projects to your training experience? *

| | 1 | 2 | 3 | 4 | 5 | 6 | 7 | |
|---|---|---|---|---|---|---|---|---|
| Not at all valuable | ○ | ○ | ○ | ○ | ○ | ○ | ○ | Extremely valuable |

How valuable is/was writing code to your training experience? *

| | 1 | 2 | 3 | 4 | 5 | 6 | 7 | |
|---|---|---|---|---|---|---|---|---|
| Not at all valuable | ○ | ○ | ○ | ○ | ○ | ○ | ○ | Extremely valuable |

How valuable is/was working on data-science-heavy-projects to your training experience? *

| | 1 | 2 | 3 | 4 | 5 | 6 | 7 | |
|---|---|---|---|---|---|---|---|---|
| Not at all valuable | ○ | ○ | ○ | ○ | ○ | ○ | ○ | Extremely valuable |

How valuable is/was writing papers to your training experience? *

| | 1 | 2 | 3 | 4 | 5 | 6 | 7 | |
|---|---|---|---|---|---|---|---|---|
| Not at all valuable | ○ | ○ | ○ | ○ | ○ | ○ | ○ | Extremely valuable |

How valuable is/was training with/from permanent staff members (staff scientists/SWEs) to your training experience? *

| | 1 | 2 | 3 | 4 | 5 | 6 | 7 | |
|---|---|---|---|---|---|---|---|---|
| Not at all valuable | ○ | ○ | ○ | ○ | ○ | ○ | ○ | Extremely valuable |

How valuable is/was training with/from other postdoc program members to your training experience? *

| | 1 | 2 | 3 | 4 | 5 | 6 | 7 | |
|---|---|---|---|---|---|---|---|---|
| Not at all valuable | ○ | ○ | ○ | ○ | ○ | ○ | ○ | Extremely valuable |

How valuable is/was interaction with collaborators to your training experience? *

| | 1 | 2 | 3 | 4 | 5 | 6 | 7 | |
|---|---|---|---|---|---|---|---|---|
| Not at all valuable | ○ | ○ | ○ | ○ | ○ | ○ | ○ | Extremely valuable |

How valuable is/was explicit and/or implicit training on project management skills to your training experience? *

| | 1 | 2 | 3 | 4 | 5 | 6 | 7 | |
|---|---|---|---|---|---|---|---|---|
| Not at all valuable | ○ | ○ | ○ | ○ | ○ | ○ | ○ | Extremely valuable |

How valuable is/was explicit and/or implicit training on communication skills to your training experience? *

| | 1 | 2 | 3 | 4 | 5 | 6 | 7 | |
|---|---|---|---|---|---|---|---|---|
| Not at all valuable | ○ | ○ | ○ | ○ | ○ | ○ | ○ | Extremely valuable |

Rank these aspects of the program in terms of most to least important. *

| | 1st | 2nd | 3rd | 4th | 5th | 6th | 7th | 8th | 9t |
|---|---|---|---|---|---|---|---|---|---|
| Curriculum | ○ | ○ | ○ | ○ | ○ | ○ | ○ | ○ | ○ |
| Workshops | ○ | ○ | ○ | ○ | ○ | ○ | ○ | ○ | ○ |
| Forum | ○ | ○ | ○ | ○ | ○ | ○ | ○ | ○ | ○ |
| Analysis projects | ○ | ○ | ○ | ○ | ○ | ○ | ○ | ○ | ○ |
| Code | ○ | ○ | ○ | ○ | ○ | ○ | ○ | ○ | ○ |
| Data science | ○ | ○ | ○ | ○ | ○ | ○ | ○ | ○ | ○ |
| Papers | ○ | ○ | ○ | ○ | ○ | ○ | ○ | ○ | ○ |
| Staff training | ○ | ○ | ○ | ○ | ○ | ○ | ○ | ○ | ○ |
| Peer training | ○ | ○ | ○ | ○ | ○ | ○ | ○ | ○ | ○ |
| Collaborator interaction | ○ | ○ | ○ | ○ | ○ | ○ | ○ | ○ | ○ |
| Project management | ○ | ○ | ○ | ○ | ○ | ○ | ○ | ○ | ○ |
| Communication skills | ○ | ○ | ○ | ○ | ○ | ○ | ○ | ○ | ○ |

Next

Clear form

Never submit passwords through Google Forms.

### Postdoctoral Training Program Feedback form

Not shared

* Indicates required question

About the program - current members

What in your opinion is the ideal length of this program? *

- [ ] 6 mo
- [ ] 6 - 12 mo
- [ ] 12 - 18 mo
- [ ] 18 - 24 mo
- [ ] 24 - 30 mo
- [ ] 30 - 36 mo

What were your career goals before you started the program and how (if at all) have your career goals changed since joining the program?

Your answer

Back Next Clear form

Never submit passwords through Google Forms.

This form was created inside of Broad Institute of MIT and Harvard. Report Abuse

Google Forms

### Postdoctoral Training Program Feedback form

Not shared

* Indicates required question

About the program - past members

What in your opinion is the ideal length of this program? *

- [ ] 6 mo
- [ ] 6 - 12 mo
- [ ] 12 - 18 mo
- [ ] 18 - 24 mo
- [ ] 24 - 30 mo
- [ ] 30 - 36 mo

What were your career goals before you started the program and how (if at all) did they change during or since the program?

Your answer

Based purely on training, how was the program length? *

- ◯ Significantly too short
- ◯ About right
- ◯ Significantly too long

When also considering the impact on your personal life that being in program entails (e.g. salary, location requirement) , how was the program length? *

- ◯ Significantly too short
- ◯ About right
- ◯ Significantly too long

When looking back, do you think joining a training program was the right choice for you? *

- ◯ Yes
- ◯ No
- ◯ Maybe
- ◯ Other:

Back Next Clear form

Never submit passwords through Google Forms.

This form was created inside of Broad Institute of MIT and Harvard. Report Abuse

Google Forms

### Postdoctoral Training Program Feedback form

Not shared

Open questions

What aspects not already mentioned do you think were very important to your training experience? Feel free to provide multiple answers.

Your answer

What aspects do you think were missing from your training experience? Feel free to provide multiple answers.

Your answer

Anything else you'd want to add that you're ok sharing as part of the paper?

Your answer

Anything else you'd want to add that you want to stay within the platform? As a reminder, if you want to send anything to Beth's eyes only, you can do it here.

Your answer
